# TMPRSS2 and RNA-dependent RNA polymerase are effective targets of therapeutic intervention for treatment of COVID-19 caused by SARS-CoV-2 variants (B.1.1.7 and B.1.351)

**DOI:** 10.1101/2021.04.06.438540

**Authors:** Jihye Lee, JinAh Lee, Hyeon Ju Kim, Meehyun Ko, Youngmee Jee, Seungtaek Kim

## Abstract

SARS-CoV-2 is a causative agent of COVID-19 pandemic and the development of therapeutic interventions is urgently needed. So far, monoclonal antibodies and drug repositioning are the main methods for drug development and this effort was partially successful. Since the beginning of COVID-19 pandemic, the emergence of SARS-CoV-2 variants has been reported in many parts of the world and the main concern is whether the current vaccines and therapeutics are still effective against these variant viruses. The viral entry and viral RNA-dependent RNA polymerase (RdRp) are the main targets of current drug development, thus the inhibitory effects of TMPRSS2 and RdRp inhibitors were compared among the early SARS-CoV-2 isolate (lineage A) and the two recent variants (lineage B.1.1.7 and lineage B.1.351) identified in the UK and South Africa, respectively. Our in vitro analysis of viral replication showed that the drugs targeting TMPRSS2 and RdRp are equally effective against the two variants of concern.

## Introduction

COVID-19 is an emerging infectious disease caused by a novel coronavirus, severe acute respiratory syndrome coronavirus 2 (SARS-CoV-2) (1) and it was declared as a pandemic by the World Health Organization (WHO) on March 11, 2020. In order to address this unprecedented global challenge, intensive investigations have been simultaneously conducted by global scientific communities and industries to develop diagnostic tools, vaccines, and therapeutics. Remarkably, within ten months after release of the SARS-CoV-2 genome sequence, a couple of vaccines were successfully developed and are now being used for vaccination of people after emergency use authorization (EUA). Drug development was also partially successful, especially in the development of monoclonal antibodies (2)(3). Notably, the vaccines and monoclonal antibodies currently being used are heavily dependent on the structure and sequence of viral Spike protein, which is a surface glycoprotein responsible for virus entry by interacting with the host receptor, angiotensin-converting enzyme 2 (ACE2). Thus, if there is any mutation in this protein, it is likely to affect the efficacy of both vaccines and antibodies.

Since the beginning of COVID-19 pandemic, variants of SARS-CoV-2 have been reported in many parts of the world and the recent variants identified in the UK (lineage B.1.1.7), South Africa (lineage B.1.351), and Brazil (lineage P.1) are of particular concern due to multiple mutations in the Spike gene (Figure 1) (4)(5). Indeed, several results are being published, which demonstrated reduced neutralization capacity of convalescent plasma, vaccine sera, and monoclonal antibodies against these variants (6)(7)(8)(9).

**Figure 1.**
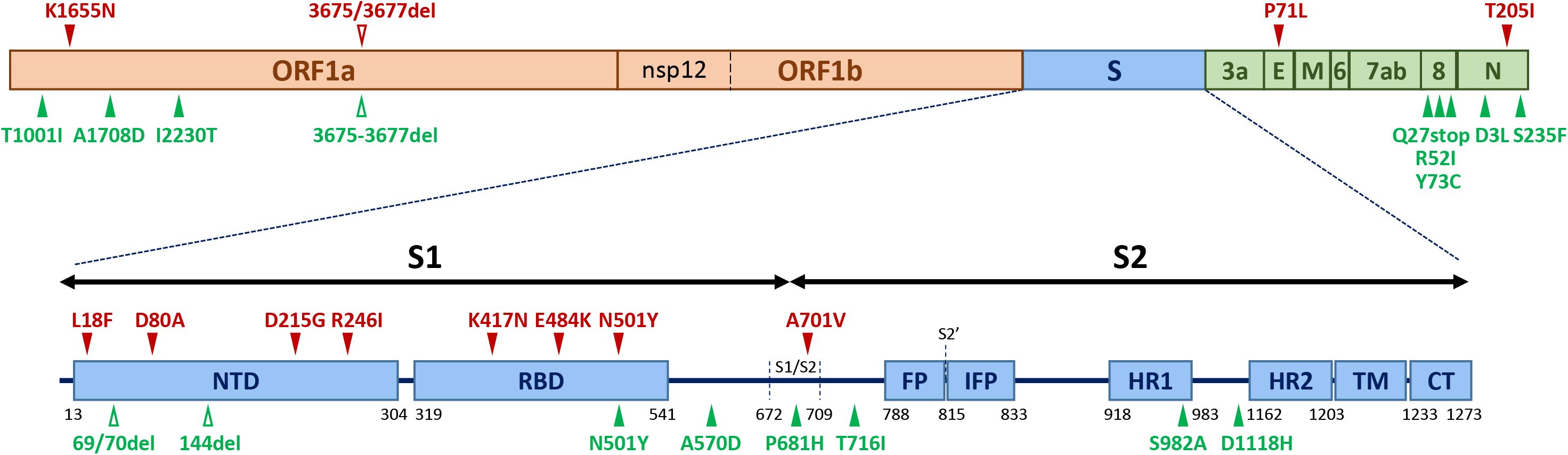
Schematic illustration of single-nucleotide polymorphisms (SNPs) in SARS-CoV-2 variants. Three SARS-CoV-2 lineages were used in this study; lineage A (an early SARS-CoV-2 isolate), lineage B.1.1.7 (identified in the UK), and lineage B.1.351 (identified in South Africa). SNPs that are observed in B.1.351 compared to the early isolate are noted in red above the diagram. SNPs observed in B.1.1.7 compared to the early isolate are noted in green below the diagram. NTD (N-terminal domain); RBD (receptor-binding domain); FP (fusion peptide); IFP (internal fusion peptide); HR1 (heptad repeat 1); HR2 (heptad repeat 2); TM (transmembrane anchor); CT (cytoplasmic tail)

In addition to monoclonal antibodies, small molecule inhibitors are also being developed as potential antiviral agents. Targets of such small molecule inhibitors are often transmembrane serine protease 2 (TMPRSS2) (10)(11)(12)(13) and viral RNA-dependent RNA polymerase (RdRp) (14)(15). TMPRSS2 is known to possess serine protease activity, which primes the viral Spike protein for fusion between the viral membrane and the host cell membrane prior to the release of viral genome into the cytoplasm. Camostat and nafamostat are representative drug candidates as TMPRSS2 inhibitors and currently being tested in several phase 2 and 3 clinical trials in many countries. On the other hand, RdRp is a target of remdesivir, which is the first approved drug for treatment of COVID-19 patients (16).

In this study, we investigated whether the antiviral drug candidates targeting TMPRSS2 and RdRp are still effective against the recent SARS-CoV-2 variants of concern by assessing in vitro viral replication capacity after drug treatment.

## Results and Discussion

The alignment of SARS-CoV-2 amino acid sequences of two lineages (B.1.1.7 and B.1.351) identified numerous changes compared to the sequence of the early SARS-CoV-2 isolate (lineage A) and several of them were located in the Spike protein (Figure 1) while no change was observed in the NSP12 amino acid sequence which possesses an RdRp activity.

In order to compare the drug susceptibilities against the three lineages of SARS-CoV-2, both Vero and Calu-3 cells were used for virus infection and drug treatment. Drugs were added to the cells prior to the virus infection. The cells were fixed at 24 h post infection and scored by immunofluorescence analysis with an antibody specific for the viral N protein. The microscopic images of both viral N protein and cell nuclei were analyzed using the Columbus software and the dose-response curve (DRC) for each drug and variant was generated (Figures 2 and 3).

**Figure 2.**
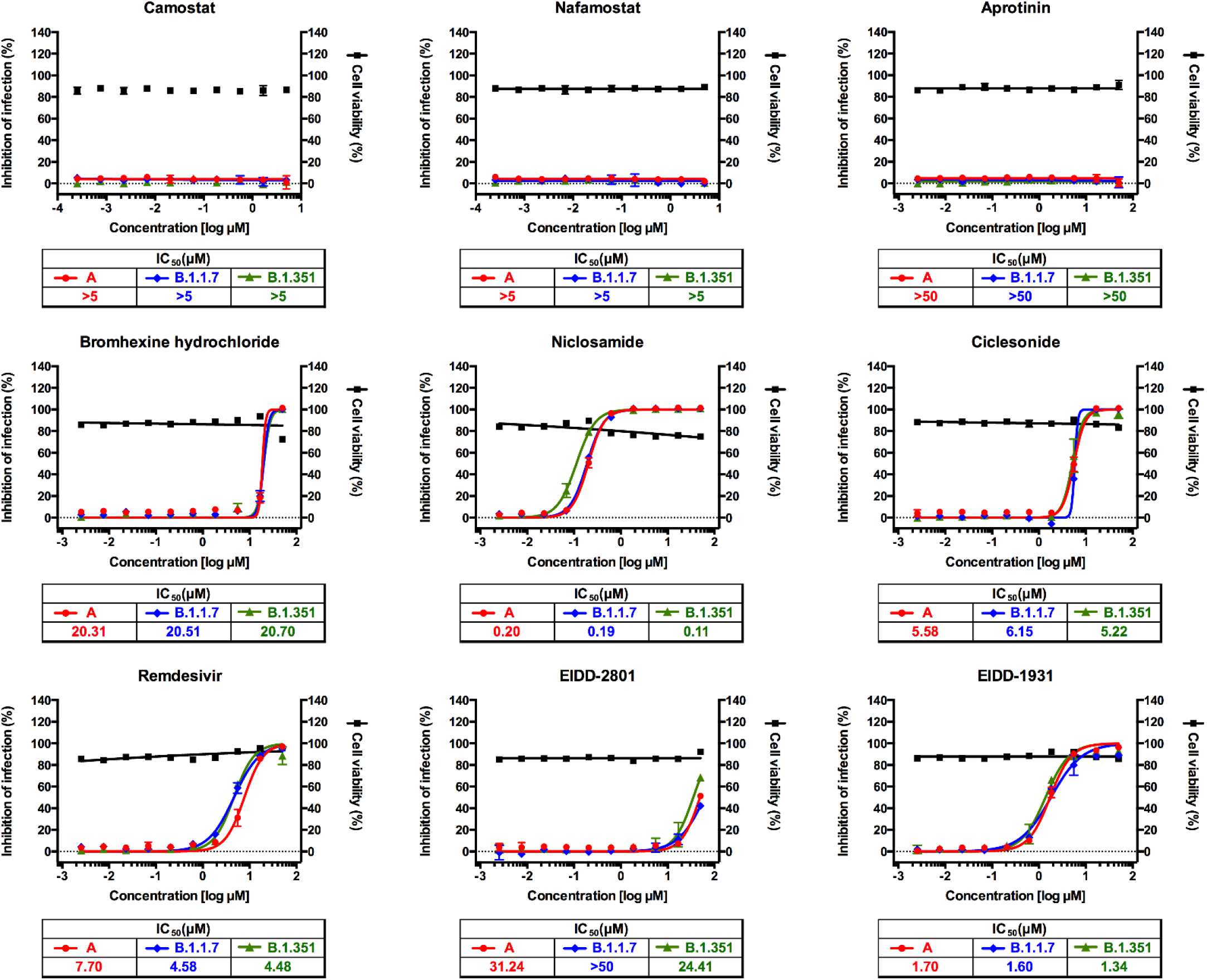
Dose-response curve analysis in Vero cells for the 9 drugs that were tested in this study. The red circles (lineage A), blue diamonds (lineage B.1.1.7), and green triangles (lineage B.1.351) represent inhibition of SARS-CoV-2 infection (%) in the presence of increasing concentrations of each drug, and the black squares represent cell viability (%). Means ± SD were calculated from duplicate experiments.

**Figure 3.**
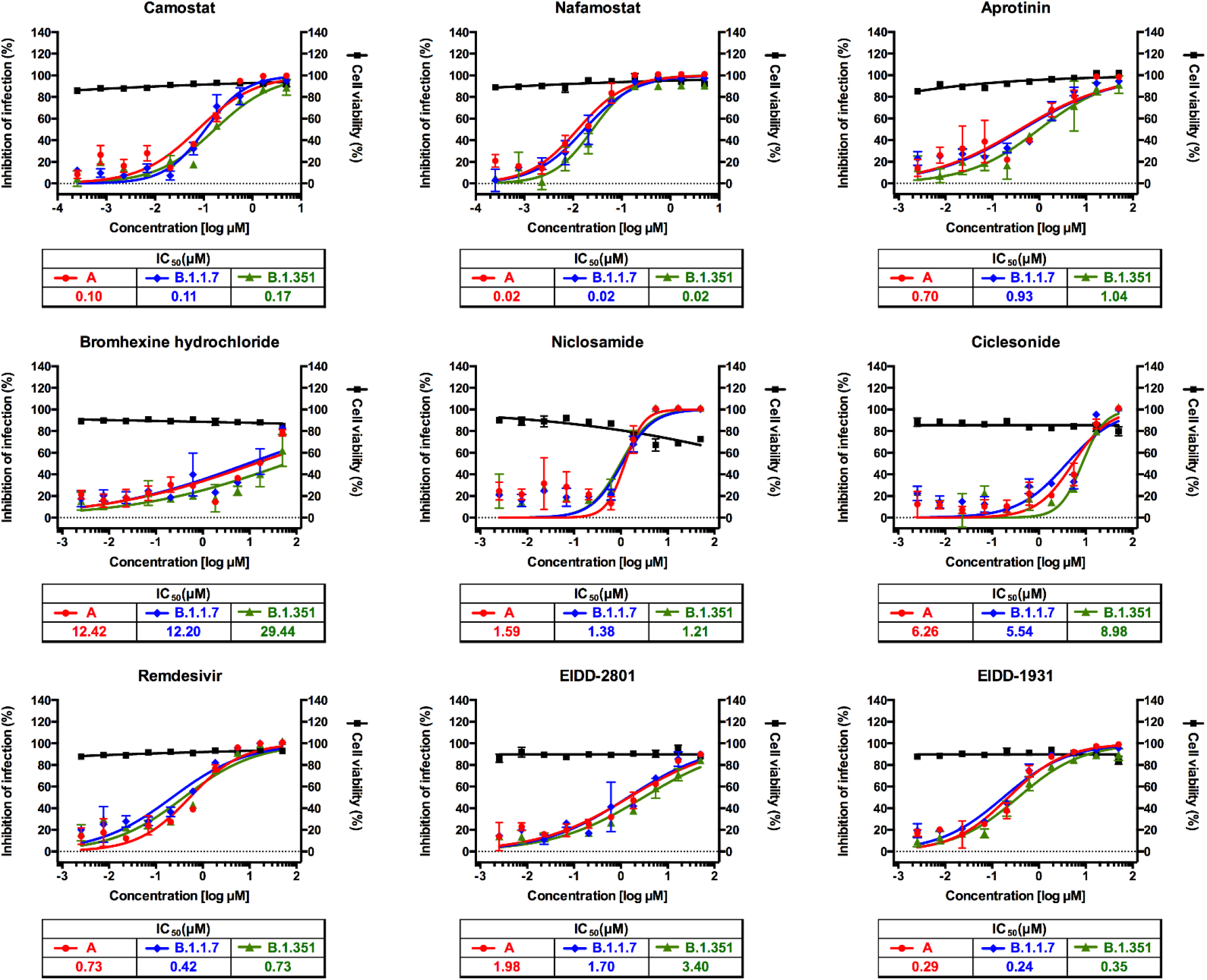
Dose-response curve analysis in Calu-3 cells for the 9 drugs that were tested in this study. The red circles (lineage A), blue diamonds (lineage B.1.1.7), and green triangles (lineage B.1.351) represent inhibition of SARS-CoV-2 infection (%) in the presence of increasing concentrations of each drug, and the black squares represent cell viability (%). Means ± SD were calculated from duplicate experiments.

We tested four different TMPRSS2 inhibitors (camostat, nafamostat, aprotinin, and bromhexine) (17), two RdRp inhibitors (remdesivir, EIDD-2801 (molnupiravir), and EIDD-1931 (an active form of EIDD-2801)) (14)(15), and others (niclosamide and ciclesonide) that we had identified in our earlier drug repositioning study (13)(18). The antiviral drug efficacy of each drug was compared among the three lineages of SARS-CoV-2; A (an early SARS-CoV-2 isolate), B.1.1.7 (identified in the UK) and B.1.351 (identified in South Africa).

While TMPRSS2 inhibitors did not show any antiviral effect in Vero cells as reported previously (Figure 2) (13), they were very effective in suppressing viral replication in Calu-3 cells, perhaps due to the abundant TMPRSS2 expression in this cell line (19) without substantial differences in drug efficacy among the three lineages of SARS-CoV-2 (Figure 3). TMPRSS2 cleaves the Spike protein at the S2’ cleavage site and no sequence change was observed at or near this site in the two recent variants (B.1.1.7 and B.1.351) compared to the sequence of the early SARS-CoV-2 isolate (lineage A) (Figure 1). Perhaps, the conserved sequence at this region could account for the similar drug efficacy among the three lineages.

The amino acid sequence of NSP12 was also well conserved among the three lineages of SARS-CoV-2 (Figure 1) and we did not find any substantial differences among them with regard to drug efficacy of the two representative RdRp inhibitors (remdesivir and molnupiravir) (Figures 2 and 3). Both remdesivir and molnupiravir are nucleoside analogs but the two drugs differ from each other in that remdesivir works as a chain terminator but molnupiravir induces mutations during viral RNA replication. Molnupiravir (EIDD-2801) is a prodrug of β-D-N^4^-hydroxycytidine (EIDD-1931) and it has well-known broad-spectrum antiviral activity against various RNA viruses (20)(21)(22)(23). Since this drug is orally available, it could be easily administered for the patients even with mild COVID-19 if it is successfully developed. Currently, phase 2 and 3 clinical trials are being conducted globally for this new drug candidate.

Finally, we assessed the antiviral drug efficacy of niclosamide and ciclesonide and no substantial differences in drug efficacy was observed among the three lineages (Figures 2 and 3). This result suggests that the potential targets of these drugs lie outside of the substituted amino acids in the two variants. Currently, niclosamide and ciclesonide are being tested in several clinical trials to assess antiviral efficacy against SARS-CoV-2 infection.

Most of monoclonal antibodies, convalescent plasma, and vaccines that are being used for treatment or prevention of COVID-19 were developed to target the viral Spike protein, specifically, the receptor-binding domain. While this protein is abundant and more immunogenic than the other viral proteins, it is also the place where many mutations occur (e.g., N501Y, E484K, K417N) due to potential viral adaptations and various selective pressures, etc. Of these mutations, some are known to substantially reduce neutralization capacity of monoclonal antibodies, convalescent plasma, and vaccine sera. Hence, it is very important to develop therapeutics targeting viral or host factors other than the Spike protein in order to address potential resistance issues caused by the Spike mutations.

In summary, we analyzed efficacy of potential drug candidates (i.e., TMPRSS2 inhibitors, RdRp inhibitors and others) against the recent SARS-CoV-2 variants of concern and we found that all of them were equally effective in suppressing replication of B.1.1.7 and B.1.351 variants compared to the early SARS-CoV-2 isolate. The results from this study would help develop therapeutic interventions specifically targeting TMPRSS2, RdRp or other viral and host factors.

## Materials and Methods

### Virus and cells

Vero and Vero E6 cells were obtained from the American Type Culture Collection (ATCC CCL-81 and C1008, respectively) and maintained at 37°C with 5% CO_2_ in Dulbecco’s modified Eagle’s medium (DMEM; Welgene), supplemented with 10% heat-inactivated fetal bovine serum (FBS) and 2% antibiotic-antimycotic solution (Gibco). Calu‐3 used in this study is a clonal isolate, which shows higher growth rate compared with the parental Calu‐3 obtained from the American Type Culture Collection (ATCC, HTB‐55). Calu‐3 was maintained at 37°C with 5% CO_2_ in Eagle’s Minimum Essential Medium (EMEM, ATCC) supplemented with 20% heat‐inactivated fetal bovine serum (FBS), 1% MEM-Non-Essential Amino Acid solution (Gibco) and 2% antibiotic‐antimycotic solution (Gibco). Three lineages of SARS‐CoV‐2 were provided by Korea Disease Control and Prevention Agency (KDCA), and were propagated in Vero E6 cells. Each lineage is noted as lineage A (an early SARS-CoV-2 isolate) (hCoV-19/Korea/KCDC03/2020), lineage B.1.1.7 (hCoV-19/Korea/KDCA51463/2021), and lineage B.1.351 (hCoV-19/Korea/KDCA55905/2021) in this study. Viral titers were determined by plaque assays in Vero cells. All experiments using SARS‐CoV‐2 were performed at Institut Pasteur Korea in compliance with the guidelines of the KNIH, using enhanced biosafety level 3 (BSL‐3) containment procedures in laboratories approved for use by the KDCA.

### Reagents

All compounds except for ciclesonide and EIDD-1931 were purchased from MedChemExpress (Monmouth Junction, NJ). Ciclesonide and EIDD-1931 were purchased from Cayman Chemical (AnnArbor, MI). Stock solution was dissolved in dimethyl sulfoxide (DMSO) at 10mM concentration. Anti‐SARS‐CoV‐2 N protein antibody was purchased from Sino Biological Inc (Beijing, China). Alexa Fluor 488 goat anti‐rabbit IgG (H + L) secondary antibody and Hoechst 33342 were purchased from Molecular Probes. Paraformaldehyde (PFA) (32% aqueous solution) and normal goat serum were purchased from Electron Microscopy Sciences (Hatfield, PA) and Vector Laboratories, Inc (Burlingame, CA), respectively.

### Dose-response curve (DRC) analysis

Vero cells were seeded at 1.0 × 10^4^cells per well with Dulbecco’s modified Eagle’s medium (DMEM; Welgene) supplemented with 2% heat-inactivated fetal bovine serum (FBS) and 2% antibiotic-antimycotic solution (Gibco) in a black, 384‐well, μClear plates (Greiner Bio‐One) 24 hours before the experiment. Calu-3 cells were seeded at 2.0 × 10^4^cells per well with Eagle’s Minimum Essential Medium (EMEM, ATCC) supplemented with 20% heat-inactivated fetal bovine serum (FBS), 1% MEM-Non-Essential Amino Acid solution (Gibco) and 2% antibiotic-antimycotic solution (Gibco) in a black, 384‐well, μClear plates (Greiner Bio‐One) 24 hours before the experiment. Ten‐point DRCs were generated with three-fold dilutions, with compound concentrations ranging from 0.0025 to 50 μM. Only nafamostat and camostat used a top concentration of 5 μM instead of 50 μM, thus concentrations ranged from 0.00025 to 50 μM. For viral infection, plates were transferred into the BSL‐3 containment facility and SARS‐CoV‐2 was added at a multiplicity of infection of 0.008 for Vero cells and 0.2 for Calu-3 cells. The plates were incubated at 37ºC for 24 hours. The cells were fixed at 24 hours post infection (hpi) with 4% paraformaldehyde (PFA), permeabilized with 0.25% Triton X-100 solution. Anti-SARS-CoV-2 nucleocapsid (N) primary antibody, 488-conjugated goat anti-rabbit IgG secondary antibody and Hoechst 33342 were treated to the cells for immunofluorescence. The images acquired with Operetta high-throughput imaging device (Perkin Elmer) were analyzed using the Columbus software (Perkin Elmer) to quantify cell numbers and infection ratios. Antiviral activity was normalized to infection control (0.5% DMSO) in each assay plate. Cell viability was measured by counting nuclei in each well and normalizing it to the mock control. DRCs were generated using Prism7 software (GraphPad). IC_50_ values were calculated using nonlinear regression analysis – log[inhibitor] vs. response – Variable slope (four parameters). All IC_50_ and CC_50_ values were measured in duplicates.

## Acknowledgements

The pathogen resources (NCCP43326, NCCP43381, and NCCP43382) for this study were provided by the National Culture Collection for Pathogens. This work was supported by the National Research Foundation of Korea (NRF) grant funded by the Korean government (MSIT) (NRF-2017M3A9G6068245, NRF-2020M3E9A1041756, and NRF-2020M3A9I2081692).

## Notes

### Competing Interest Statement

The authors have declared no competing interest.

